# Impacts of heat stress and its mitigation by capsaicin in health status and digestive enzymes in house sparrows (*Passer domesticus*)

**DOI:** 10.1101/2023.04.17.537187

**Authors:** Julia Cacace, Guido Fernández Marinone, Fabricio Damián Cid, Juan Gabriel Chediack

## Abstract

Heatwaves and warm spells at global level, product of climate change, causes alterations on homeostasis in animals (heat stress), so they must respond to these changes in order to survive. The response involves several physiological changes, such as releasing of glucocorticoids and catecholamines, to restore homeostasis. Due the lack of knowledge on this subject in wild birds, the objective of the present work was study the effect of heat stress on body condition and digestive system in house sparrows (*Passer domesticus*), as well as a possible role of capsaicin as a dietary additive in mitigating heat stress. In this work, we measure heterophils/lymphocytes ratio (as proxy of stress), body mass, hematocrit, uric acid and digestive enzymes (intestinal and pancreatic) under stress conditions at 32°±2 °C and under normal conditions at 22°±2 °C. In addition, we evaluate the effect of capsaicin in both situations (heat stress at 32°C and normal condition 22°C). We found an increase of H/L ratio and uric acid in plasma under heat stress, and decrease of H/L ratio with capsaicin on animals exposed to heat stress. Although a loss of intestinal mass was observed in animals exposed at 32°C, digestive enzyme activity does not change under heat stress or under capsaicin administration. Improving knowledge in this field is relevant at the level of animal nutrition and veterinary medicine, reducing the stress of wild birds in captivity and improving dietary mixes for future global warming scenarios.

## 1. Introduction

The increase of frequency, intensity and duration of heatwaves and warm spells at global level are climatic phenomena produced by climate changes (Perkins et al, 2012; Perkins-Kirkpatrick and Lewis, 2020). This environmental instability causes in the animals an alteration of their homeostasis (animal stress) and they must adapt faster to these changes in order to survive (Breuner, 2003). The stress response involves several physiological responses governed primarily by activation of the sympathetic-adrenal (SA) axis and the hypothalamic-pituitary-adrenal (HPA) axis, releasing hormones (glucocorticoids and catecholamines) into the bloodstream (Koolhaas et al., 1999; Wingfield and Kitaysky, 2002). In birds, the main glucocorticoid hormone released by stress is corticosterone (Siegel, 1994; Hess, 2002; Sheriff et al, 2011) and one of its effects is to increase the H/L ratio (Gross and Siegel, 1983; Davis et al, 2008; Chediack et al, 2022). An increase in corticosterone (CORT) has been observed in wild birds under different stress events, such as fasting (Chediack et al, 2022), season or reproductive stage (Romero et al, 2006), and heat stress (Bruijn and Romero, 2011; Xie et al, 2017; Newberry and Swanson, 2018). But there is a lack of knowledge in wild animals, particularly in birds, regarding heat stress on blood parameters. Also, it is not known whether the digestive process, such as intestinal enzyme activity, could be affected. Most heat stress studies come from poultry. On the other hand, there are controversial information about its effect on some blood parameters, such as uric acid and triglycerides (Sughiarto et al, 2017; Bueno et al, 2017; Bogin et al, 1996, Xie et al, 2015; John and George, 1977).

In birds, it has been well documented that exposure to stressors (fasting, disturbing events, heat stress, etc.) elevates CORT blood concentration, and its consequence is an increase of heterophils blood concentration, while lymphocytes are diminished (Post et al, 2003; Shini et al, 2008, Chediack et al, 2022). A study shows that glucocorticoids can act to redistribute T cells from the bloodstream into organs (Ince et al, 2019). In poultry the H/L ratio increase after heat stress (Altan et al, 2000, 2003), while studies with wild desert birds show a differential response between species, with an increase of H/L ratio under heat stress conditions (Xie et al, 2017).

Studies of heat stress in the digestive system come mainly from poultry, where found different effects on gene expression in glucose transporters (SGLT1) and digestive enzymes by heat stress. (Garriga et al, 2006; Al-Zghoul et al, 2019; Goel et al, 2021). In these studies has been observed different effects on digestive enzymes activity during ontogeny and adulthood: intestinal maltase, sucrose and the pancreatic enzymes, lipase, amylase, trypsin and chymotrypsin (Osman and Tanios, 1983; Hai et al, 2000; Song et al, 2018; Wu et al, 2021). In this sense, Hao et al, (2012) observed a relationship between Hsp70 overexpression and an increase of digestive function (activity of lipase, amylase, and trypsin) in chickens. To our knowledge there are no studies in digestive physiology conducted in wild birds under heat stress.

On the other hand, the use of dietary additives as heat stress relievers has been broadly documented, for example, ascorbic acid, α-tocopherol, and folic acid produce a reduction in reactive oxygen species (ROS) in poultry (chickens and quails) exposed to heat stress (Ajakaiye et al, 2010; Gürsu et al, 2004; Ravindran, 2010). Other additives known for their anti-stress reaction include capsaicin, allicin, garlic extracts, coriander, vitamin C, and oregano essential oil (Prieto and Campo, 2010; Al-Jaff, 2011; Habibi et al, 2014; Ghazi et al, 2015; Mohamed et al, 2015). Capsaicin is a natural alkaloid and is the main active component in bell peppers/hot peppers and cayenne. It has been shown that the capsaicin administration produces a decrease in the H/L index in chickens under stress (McElroy et al, 1994; Prieto and Campo, 2010; Orndorff, 2005).

Due the lack of knowledge in wild birds, the objective of the present work was study the effect of heat stress and capsaicin as a dietary additive in mitigating heat stress: i. on body condition (body mass, organ mass); ii. on blood parameters (H/L ratio and uric acid) and iii. digestive enzymes (sucrose, maltose and aminopeptidase from intestine and trypsin and chymotrypsin from pancreas) in house sparrows (*Passer domesticus*).

As mentioned above, these studies (administration of additives in the diet of birds under heat stress) have not been documented in wild birds. Improving knowledge in this field is relevant at the level of nutritional and veterinary medicine, reducing the stress of wild birds in captivity and improving dietary mixes for future global warming scenarios.

## 2. Materials and methods

### 2.1 Animal care and housing

Adult house sparrows (*Passer domesticus,* n=24) were captured with live traps in San Luis and Mendoza province in Argentina. The birds were housed in cages (40×25×25 cm) indoors under relatively constant environmental conditions (22 ±1 °C) on a photoperiod of 12:12 h (light:dark) with food and water *ad libitum*. Food consisted of a mix of seeds (canary bird seeds and millet) supplied with vitamins and minerals, and dog food (for protein requirements). Animals were acclimated to laboratory conditions for at least one month prior to use in experiments during which birds were disturbed only for routine husbandry. Animal care and trial protocols were approved by the Animal Care and Use Committee (CICUA) of Universidad Nacional de San Luis, Argentina and were conducted in accordance with the Guide for the Care and Use of Laboratory Animals (National Research Council (U.S.) et al, 2011), protocol number N° B-69/18.

Once admitted to the animal room, the body mass of each bird was recorded, dewormed externally and internally, following the manufacturer’s instructions with a Triclorphon (Nevugón, Bayer) and then a 1 g/l piperazine deworming solution was administered in water for one week.

### 2.2 Blood extraction for biochemical and hematological parameters

Blood samples (∼200 µl total, which accounts for <10% of total blood volume; Stangel 1986) were collected by pricking the brachial vein, with a needle 25G, in a heparinized capillary tube. To avoid the influence of the circadian rhythm on the biochemical parameters fluctuation, all samples were taken at the same time (Arias et al, 2012; Chediack et al., 2022). Blood samples were centrifuged 3 min at 12,000 rpm in a hematocrit centrifuge (Cavour model VT-1224) and plasma was separated and stored in the refrigerator at 2-8 °C for blood analysis (triglycerides and uric acid). In some cases, when blood sample was hemolyzed we did not measure some parameters that could be affected by hemolysis (Bhargava et al. 2019). To determine H/L ratio, a drop of blood was smeared on a microscope slide, air dried, and stained with May-Grünwald Giemsa technique for 45 min (Gross and Siegel 1983). The H/L index was determined by the microscopic differential count on a blood smear slide stained with May-Grünwald Giemsa technique. The smears were examined under a 100x Olympus BX40 Microscope and a minimum of 100 white blood cells, heterophils (H) and lymphocytes (L), were counted to calculate the index (Fig. 2). Blood samples were taken within 3 min of removal of birds from cages to avoid the effect of corticosterone release by stress (Romero and Reed 2005).

### 2.3 Preliminary experiments

#### Effect of temperature as a stressor

To evaluate the thermal conditions that generate stress in birds, we made two experiments. In the first experiment, two groups of birds (n=7/group) were exposed to different temperatures at the same time and in contiguous rooms. A control group at 22 ±2 °C and a treatment group at 32 ±2 °C. The temperature was constantly maintained for three days. Before starting the treatment, body mass was measured and blood samples were collected for the hematocrit, and biochemical parameters (uric acid and triglycerides). H/L ratio was measured a couple of days before exposure to temperature, to avoid the effect of sampling time in the ratio. After three days, at the same time, blood samples were collected again for the same studies.

In the second experiment, one week after first one, the same group of birds was exposed to intermittent heat. A control group at 22 ±2 °C, and a treatment group at 32 ±2 °C for 11 hours (photophase) and 22 ±2 °C during night. The stoves were turned on at 8 a.m. and turned them off at 7 p.m. Before starting the treatment, body mass was measured and blood samples were collected for the hematocrit, and biochemical parameters (uric acid and triglycerides). H/L ratio was measured a couple of days before exposure to temperature, to avoid the effect of sampling time in the ratio. After three days of treatment, their body mass was measured and blood was extracted.

#### Determination of capsaicin concentration to be administered

Previous studies made by our group determine the appropriate dose of capsaicin to be used for the experiments, in mg/ml per body weight (Fernández Marinone G, 2010). The concentrations used were 15.62 x 10^-3^ mg and 31.25 x 10^-3^ mg of capsaicin (CAS 4004-86-4 Sigma) per gram of body mass. The solubility protocol was described by Turgut, 2004. The room was conditioned at 32 ±2 °C. Two stagger groups (n=6 animals in total) were used, divided by 2 birds each time. This was to avoid the addition of stress by handling the animals in the same animal room. The administration of capsaicin solution was carried out for 3 days at the same time by oral gavage (to ensure the exact amount administered). On the fourth day, their body mass was measured and blood samples were collected to perform the H/L index. Also, the hematocrit was measured and plasma samples were extracted for the determinations of the biochemical parameters.

### 2.4 Main experiment to evaluate the effect of capsaicin administration on heat stress

To generate the thermal stress, two rooms were conditioned with the same light/dark photoperiod, one room at 22 ±2 °C and the other at 32 ±2 °C. To test the effect of heat stress and its mitigation, we design four groups of birds (n=6). Group 1 (control 22): birds at 22 °C; Group 2 (treatment 22) birds at 22 °C with capsaicin; Group 3 (control 32): birds at 32 °C; Group 4 (treatment 32) birds at 32 °C with capsaicin. To avoid the influence of the circadian rhythm and to optimize the samples and data collecting-time, we worked with four birds per day. The birds were separated in 6 groups : 1 sparrow at 22 °C treated with water by oral gavage (control); 1 sparrow at 22 °C treated with capsaicin by oral gavage; 1 sparrow at 32 °C treated with water by oral gavage (control); 1 sparrow at 32 °C treated with capsaicin by oral gavage (Fig. 1). The oral gavage was made every day during 3 days. At day 4, animal surgeries were performed. Oral gavage and surgeries were carried out between 8 and 9:30 a.m. in all cases (Fig. 1).

**Figure 1.**
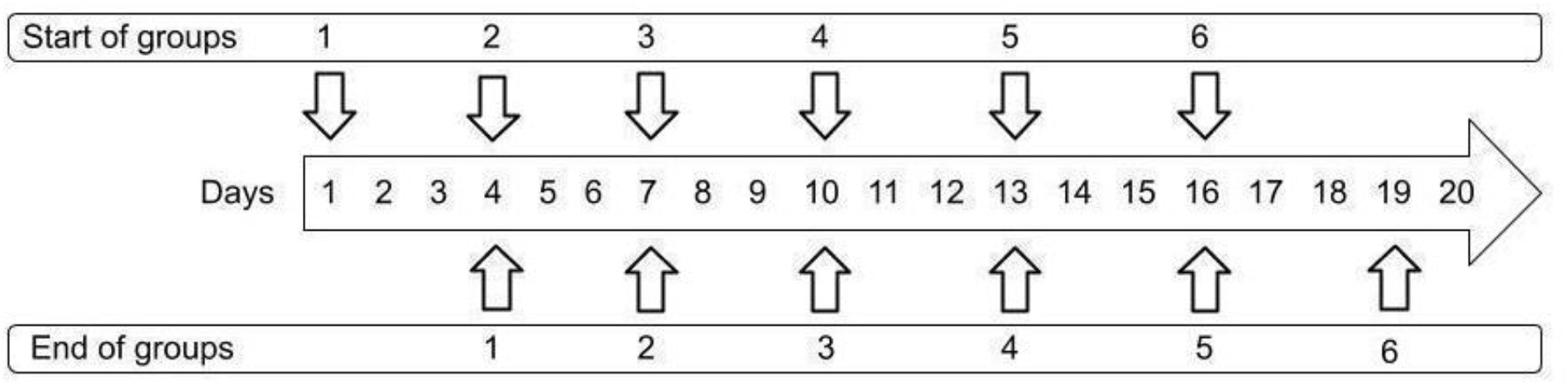
Diagram showing the timeline of the treatments.

### 2.5 Tissue extraction and homogenization

The procedures performed were approved by the Animal Care and Use Committee (CICUA) of the National University of San Luis (protocol N° B69/18). Anesthesia was induced with 40 mg.kg ketamine and 10 mg.kg xylazine diluted in saline solution and intramuscularly administered (Murphy & Fialkowski, 2001).

Prior to surgery, body mass was measured and blood was drawn. The organs were then carefully removed and placed in cold Mannitol-HBSS (pH 7) solution, on a plate with ice. The following organs were cleared of foreign tissue and weighed: small intestine, pancreas, and liver. The intestine was flushed internally with avian Ringer’s solution using a flexible cannula. After, the intestine was measured with a millimeter ruler and divided into sections (proximal, medial, and distal) taking the pylorus as a reference. These sections were weighed on an analytical balance (Denver Instrument Company A-200 DS). Intestines were kept on ice and then frozen at -80 °C until processing. Each portion was homogenized with 10 times the mass of the tissue in a HEPES-KOH solution (pH 7, 1 mM HEPES, 350 mM mannitol), with the PRO Scientific PRO200 homogenizer, in 3 intervals of no more than 10 seconds, on ice. After this, the samples were stored at -20 °C, and the activity was measured before one month. Pancreas were weighed and then frozen at -80 °C until processing. The whole pancreas was homogenized with buffer Tris/HCl 50 mM pH 8.2 with 3 mM Taurocolic acid and 0.5% Tritón X-100. Then, the samples were stored at -80 °C for subsequent analyses.

### 2.6 H/L ratio

Blood samples were taken using heparinized capillaries. A blood smear was made with a drop of blood, which was stained with May Grünwald-Giemsa. First, the smears were fixed with May-Grünwald for 5 min, then they were washed with running water. After, the smears were stained for 45 min with Giemsa diluted in water and washed with running water. Smears were observed under a light microscope at 100X to determine the ratio of two of the different types of leukocytes (lymphocytes and heterophiles) (Claver, 2009, Fig. 2). Cells in uniform fields (monolayers) were counted until 100 leukocytes were found. With the data of the count of heterophiles and lymphocytes obtained, the H/L ratio was estimated.

**Figure 2.**
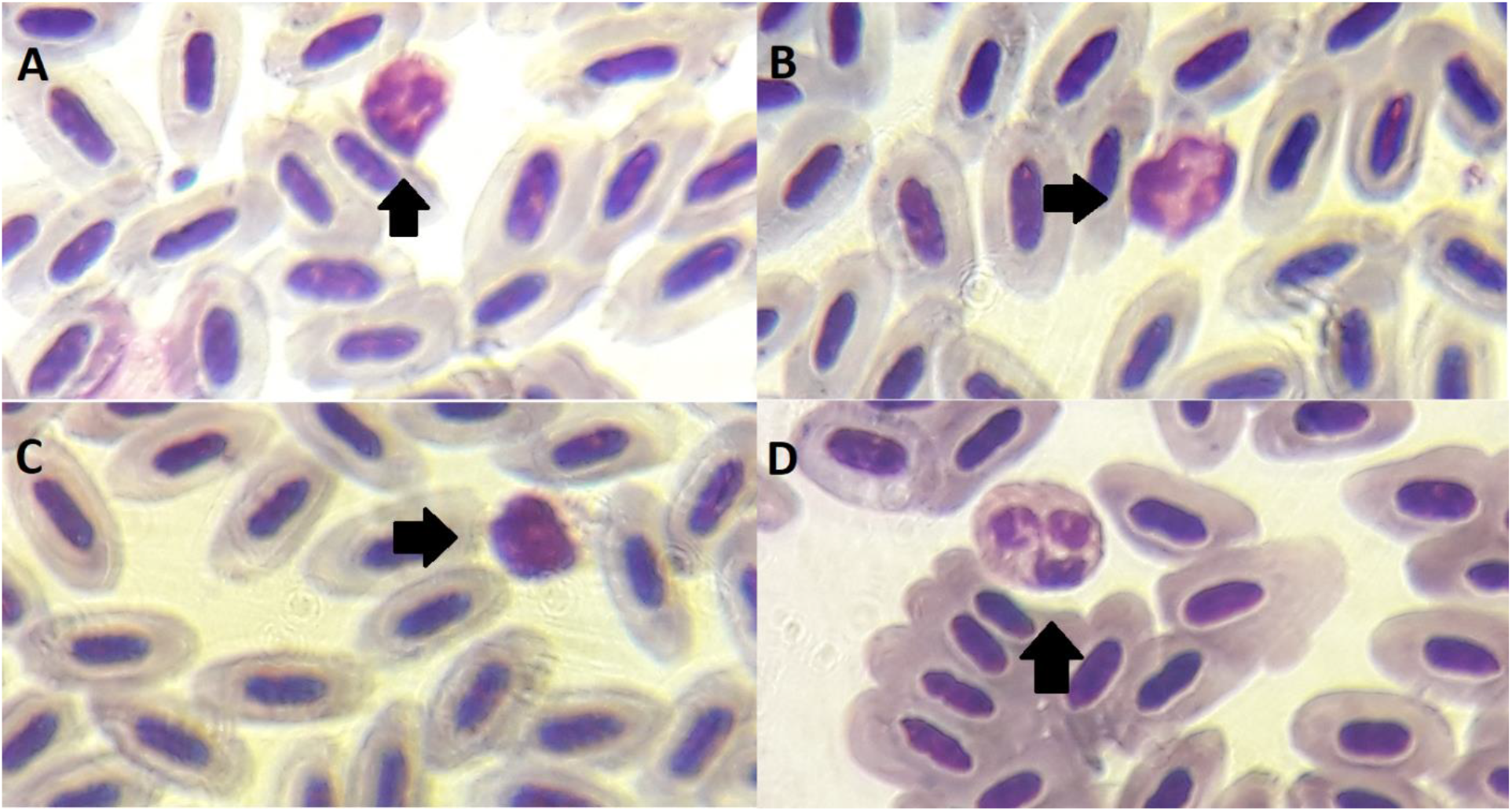
Representative light microscopic images of blood samples staining with May Grünwald Giemsa at 100X. (A-B) Mature lymphocyte. (C) Immature lymphocyte. (D) Mature heterophile.

### 2.7 Uric acid and hematocrit

#### Uric acid

Plasma samples were used to determine uric acid using commercial kits from GT Lab (GT Laboratories S.R.L, Rosario, Argentina) following standard protocols provided by the manufacturer (https://www.gtlab.com.ar). All samples were assayed in duplicate.

#### Hematocrit

Hematocrit was measured by centrifugation of heparinized capillaries for 3 min at 10,000rpm in a hematocrit centrifuge (CAVOUR model VT-1224) and subsequent reading of the percentage of red blood cells occupy in the total volume of blood. Plasma was separated and stored at -20 °C for further analysis.

### 2.8 Enzyme assays

#### Disaccharidases

To measure disaccharidase enzymes, sucrose-isomaltose (SI, EC 3.2.1.48) and maltose-glucoamilase (MG, EC 3.2.1.20) colorimetric methods of Dalqvist (1984) modified by Martínez del Rio (1990) were performed, using the enzymatic glycemia kit from Wiener Lab 40 μl of the diluted homogenates were taken and incubated, in duplicate, at 40 °C with 40 µl of substrate (56mM sucrose in 0.1 M maleate/NaOH buffer pH 6.5; 56 mM maltose in 0.1 M maleate/NaOH buffer pH 6.5) for 20 min. After incubation, the reactions were stopped byaddition of 1 ml of stop/develop solution for glucose determination (Wiener Lab). Finally, the absorbances were read at 505 nm (Spectrum Model SP-1103).

#### Intestinal peptidase

To measure aminopeptidase-N (APN, EC 3.2.11.2) a 2 mM solution of L-Ala-p-NO_2_- aniline in phosphate buffer (pH 7) was made as reaction substrate. 10 µl of the homogenate was taken and 1 ml of the substrate was added. It was incubated at 40 °C for 20 min and then a stop solution (2 M acetic acid) was added. Duplicates of each sample in addition to the blank duplicate were made. Finally, the absorbances were read at 384 nm (Spectrum Model SP- 1103).

#### Trypsin and chymotrypsin

Activity of pancreatic proteases trypsin (EC 3.4.21.4) and chymotrypsin (EC 3.4.21.1) was measured by a modification of Erlanger et al method. Pancreatic samples were homogenized, and zymogens were activated by incubation with enterokinase (1 h at 25 °C). The duration of zymogen activation was determined to insure that activities of proteases were analyzed at their maximal activation (this was checked which the duration of the zymogen activation resulted in highest activity of pancreatic proteases). Appropriately diluted aliquots were incubated with DL-BAPNA (ben-zoyl-arginine-p-nitroanilide) for trypsin or with GPNA (N-glutaryl-L-phenylalanine-p-nitroanilide) for chymotrypsin at 40 °C for 10 min. The reaction was terminated by adding 30% acetic acid (in blank samples, the acetic acid was added before the substrate). The liberated amount of p-nitroaniline was estimated by reading the absorbance at 410 nm and using a p-nitroaniline standard curve. For each enzyme, we calculated its mass-specific activity (i.e., the number of moles of product formed during 1 min incubation with a mass unit of pancreatic tissue).

### 2.9 Statistical analysis

The results are presented as the value of the mean ±1 standard error of the mean, represented in the graphs by a symbol and its respective error bars. The number of animals (n) used was at least 6 animals for each studies, except otherwise indicated.

Statistical comparisons were made using repeated measures ANOVA to examine the effect of treatments and the gut region on mass specific enzyme activities. Organ masses, as well as digestive tract measurements and biochemical parameters, were evaluated by ANOVA. Before making the comparisons, the data were tested for normality (Kolmogorov-Smirnov test) and homogeneity of variance (Levene test). In the cases where significant differences were observed, a Tukey *post hoc* test was applied. The level of significance in all tests was p ≤0.05.

## 3. Results

### Effect of temperature as a stressor

The H/L index was used to verified heat stress in house sparrows in the preliminary experiments with constant or intermittent 10 °C increase in temperature (22 °C to 32 °C), for 3 days. The value of the H/L ratio increase significantly (p<0.05) at the constant temperature treatment of 32 ±2°C, such as the intermittent temperature (Fig. 4). We chose a constant temperature treatment for the following experiments, as this treatment adds fewer other stress factors (mainly by disturbance in animal room). We did not find effect of temperature on body mass in house sparrows under heat stress (p<0.05, Fig. 3).

**Figure 3.**
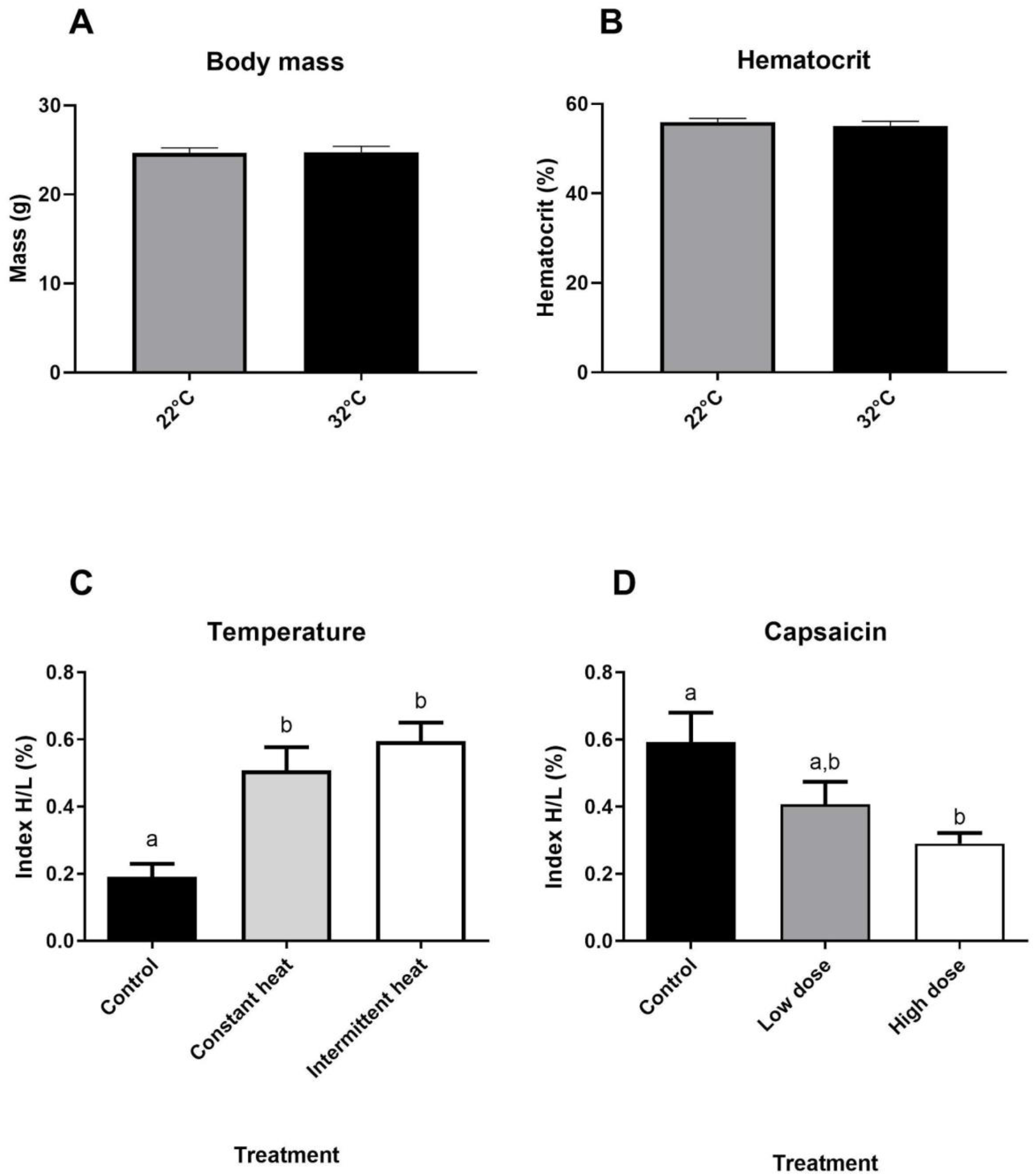
Pilot results of heat stress and capsaicin doses on H/L ratio. (A) Body mass of birds exposed to different temperatures. Bars represent the means ±S.E. (n=8). (B) Hematocrit of birds at 22±2 °C and 32 ±2 °C. Bars represent the means ±S.E. (n=7). (C) Measurement of H/L index at different temperature treatments: control, constant high temperature (Ct T), intermittent temperature (Int T). Different letters represent significant differences (p<0.05). Bars represent the means ±S.E. (n=5). (D) H/L index values at 32 ±2°C Ct T in addition to the administration of different doses of capsaicin. Different letters represent significant differences (p<0.05). Bars represent the means ±S.E. (n=6).

**Figure 4.**
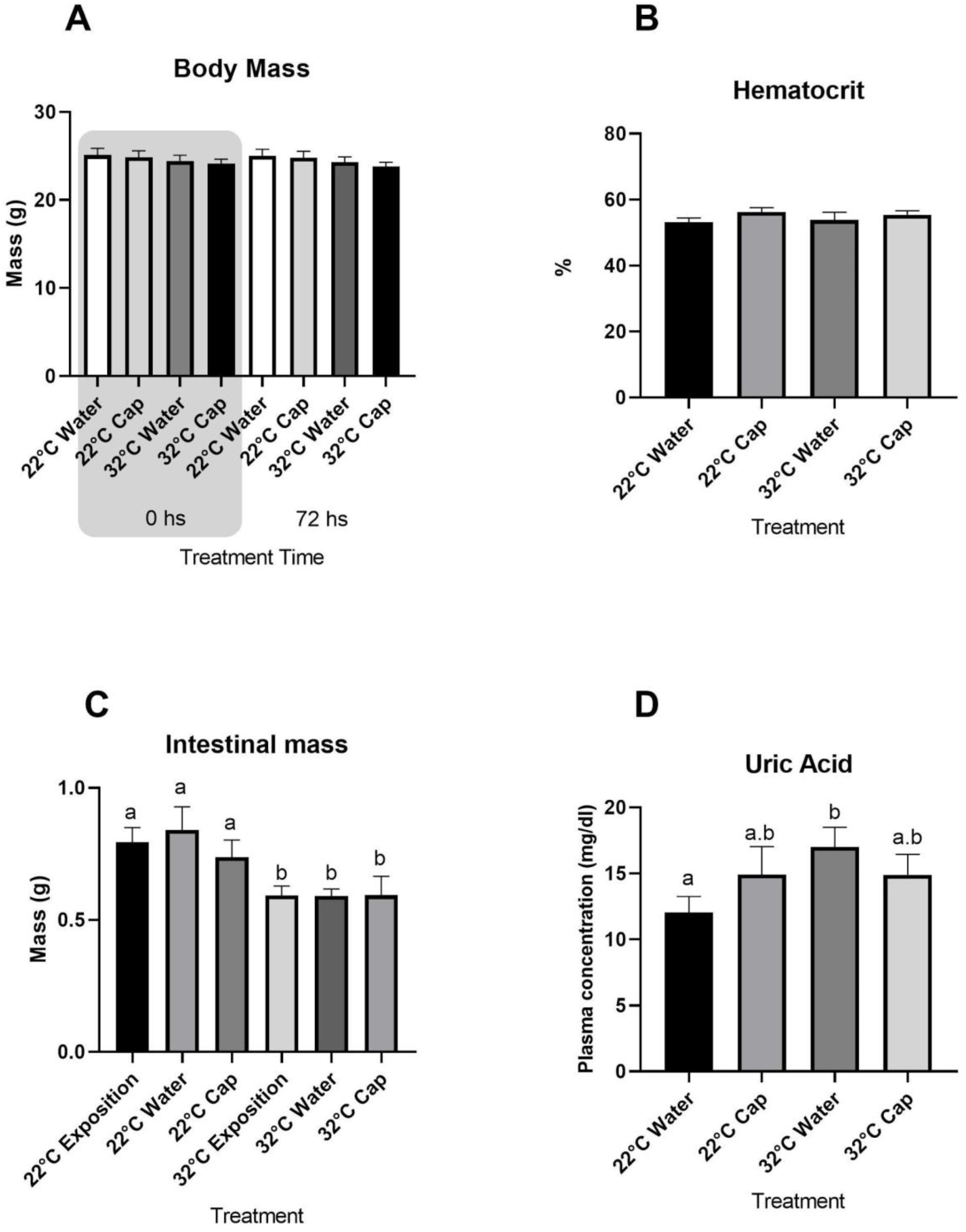
(A) Measurement of body mass in different treatments. Different letters represent significant differences (p<0.05). Bars represent the means ±S.E. (n=10 to 12). (B) Hematocrit measurement in sparrows under treatment. Different letters represent significant differences (p<0.05). Bars represent the means ±S.E. (n=6). (C) Intestinal mass after water and capsaicin treatments. Different letters represent significant differences (p<0.05). Bars represent the means ±S.E. (n=6). (D) Uric acid concentration in plasma after different treatments. Different letters represent significant differences (p<0.05). Bars represent the means ±S.E. (n=6).

### Determination of capsaicin concentration to be administered

Birds under constant heat of 32 ±2 °C for 3 days increase the H/L ratio versus control, 22 ±2 °C (Fig. 3). Capsaicin was administered in two different doses, low (15.62 x 10^-3^ mg.gr body mass^-1^ of capsaicin) and high (31.25 x 10^-3^ mg.gr body mass^-1^ capsaicin). The higher dose of capsaicin produces a significant decrease of H/L ratio respect to control (p <0.016). The values are similar of birds under temperatures of 22 ±2 °C. Low dose of capsaicin show not significantly different in the H/L ratio in comparison with the control group (p>0.216; Fig. 3).

### Main experiment to evaluate the effect of heat stress and its mitigation by capsaicin

In order to determine whether capsaicin mitigates the heat stress to which the sparrows were subjected, different parameters were analyzed.

### Body mass

No difference was observed between the masses of individuals exposed to heat stress relative to controls (p>0.5), nor control individuals exposed to capsaicin (p>0.5; Fig. 4).

### Intestine mass

Despite not observing a decrease in body mass, the intestine showed a decrease in mass with an increase in temperature from 22 °C to 32 °C (p<0.05). On the other hand, no significant differences were observed in the treatments with or without capsaicin at either the two temperatures (p>0.38; Fig. 4). An analysis of the loss of intestinal mass in relation to body mass was also performed, and a significant difference in the loss of intestinal mass was observed (Fig. 4).

### Hematocrit and uric acid

We did not observe significant differences in hematocrit values in birds under heat stress (p>0.52). Those birds under thermal stress at 32 °C had the same value as the control birds at 22 °C, all between the values 52-55% (Fig. 4).

However, heat stress increase uric acid levels in plasma (p<0.05) and capsaicin does not return to normal values of uric acid (unpaired test p = 0.2693 Fig. 4).

### Intestinal enzymes

#### Aminopeptidase

Aminopeptidase activity does not show any effect after changes in temperature or administration of capsaicin for none of the treatments (*post hoc* Tukey, p>0.1; Fig. 5). We found the same pattern (Chediack et al, 2012) of increasing activity along intestine, being lower in the proximal portion and increasing towards the distal portion (RM-ANOVA p<0.001). All activity values were analyzed as specific enzyme activity per milligram of tissue and per milligram of total protein.

**Figure 5.**
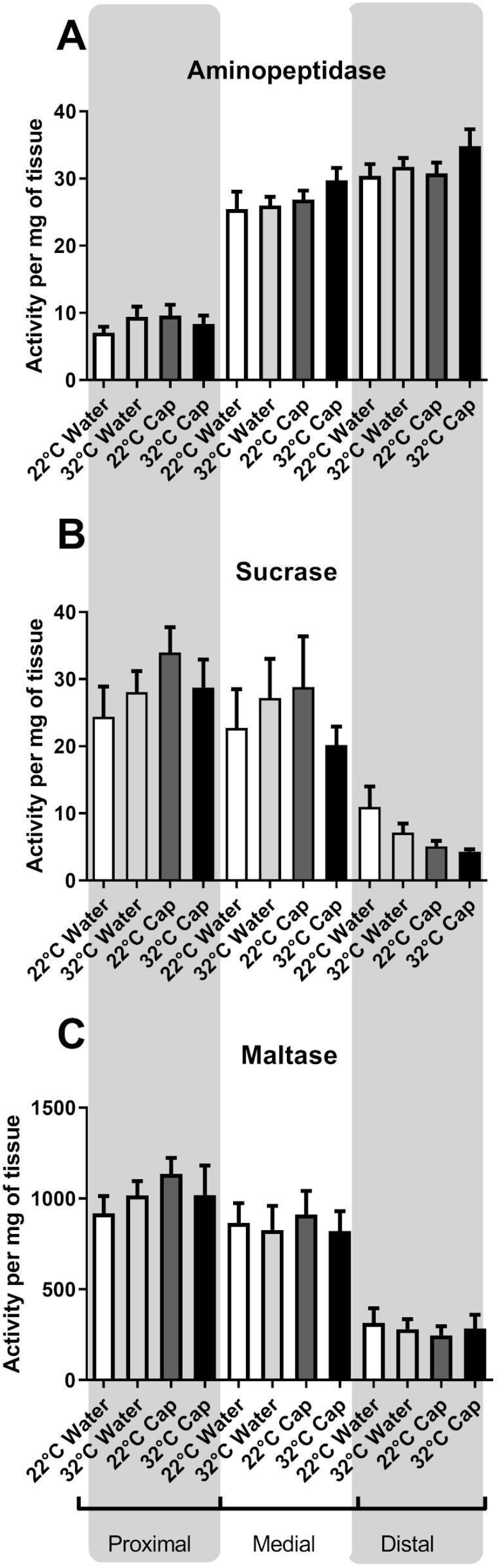
Intestinal digestive enzymes. (A) Aminopeptidase activity in the different intestinal portions under the treatments performed. Different letters represent significant differences (p<0.05). Bars represent the means ±S.E. (n=6 to 8). (B) Sucrase activity values in the different intestinal portions. Different letters represent significant differences (p<0.05). Bars represent the means ±S.E. (n=6 to 8). (C) Maltase activity values in the different intestinal portions. Different letters represent significant differences (p<0.05). Bars represent the means ±S.E. (n=6 to 8).

### Sucrase and maltase

In the case of sucrase enzyme, there is no effect of temperature or administration of capsaicin for none of the treatments (*post hoc* Tukey, p>0.1; Fig. 5). The activity pattern is similar to those found previously (Chediack et al, 2012). Highest activity values were found in the proximal portion of the small intestine and the lowest values were found in the distal portion. These values did not show significant differences in each treatment in its portion (p>0.1; Fig. 5).

Like sucrase, maltase has its highest enzyme activity values in the proximal portion and decreases to its lowest values in the distal portion of the intestine. For each treatment, in the different portions of the intestine, no significant differences were observed in their values (p>0.1; Fig. 5). Values were measured as specific enzyme activity per milligram of tissue and per milligram of total protein.

### Pancreatic enzymes: trypsin and chymotrypsin

These enzymes are secreted by the pancreas into the duodenum as zymogens. The results do not show significant differences of trypsin and chymotrypsin activities at 22 °C or 32 °C (p>0.1), and no effect in the activity by the administration of capsaicin additive (p> 0.2; Fig. 6). Values were measured as specific enzyme activity per milligram of tissue and per milligram of total protein.

**Figure 6.**
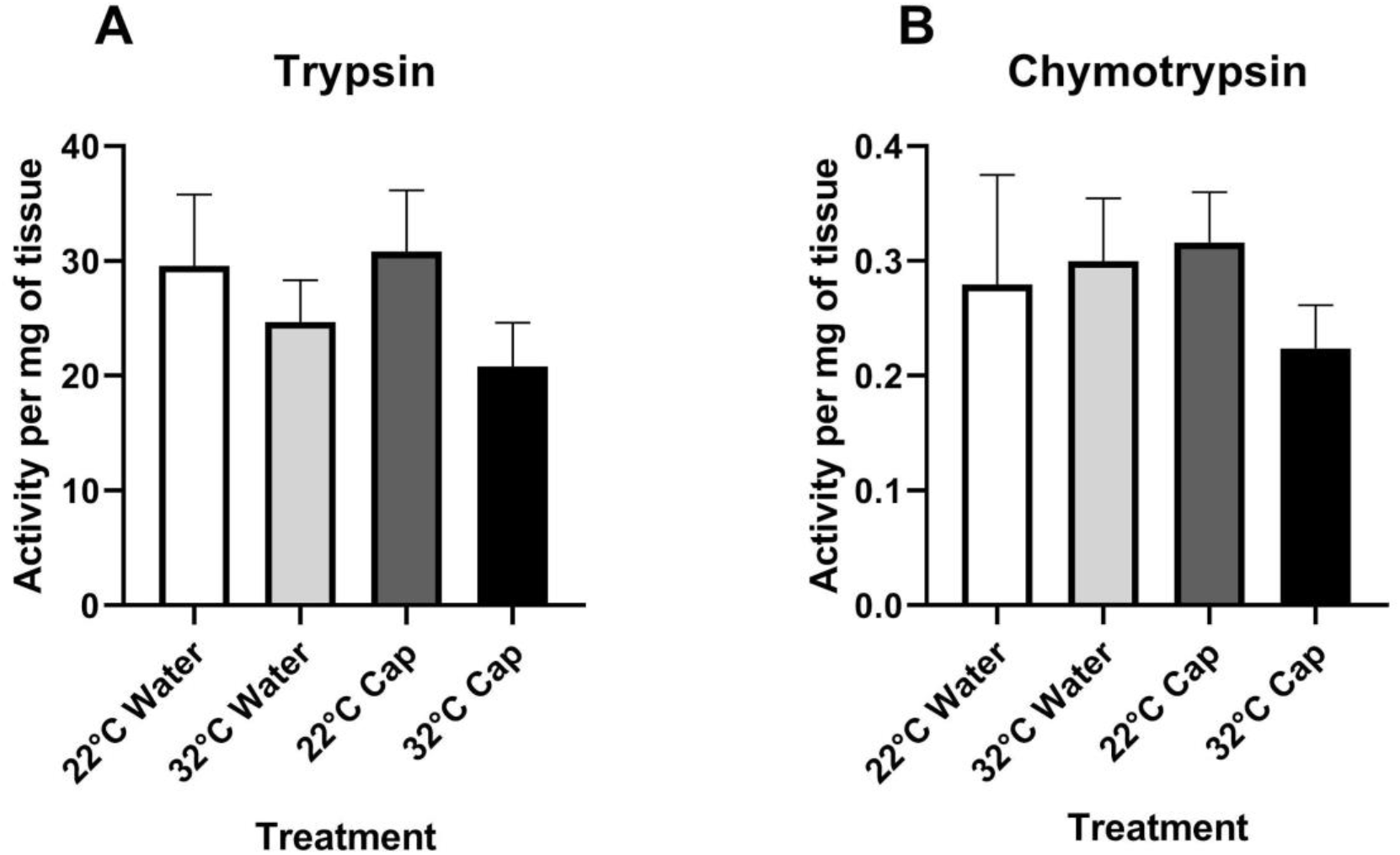
Pancreatic enzymes activity. (A) Trypsin activity values in the duodenum of the small intestine. Different letters represent significant differences (p<0.05). Bars represent the means ±S.E. (n=6). (B) Chymotrypsin activity values in the duodenum of the small intestine. Different letters represent significant differences (p<0.05). Bars represent the means ±S.E. (n=6).

## 4. Discussion

### Body mass and uric acid

Body mass and some blood parameters are useful and important as general indicators of health in individuals. Body mass, as a measure of body condition, is used in ecological and conservation studies. However, the effect of temperature body mass in wild birds is scarce and, in general, the majority comes from poultry. We did not find variations in the body mass through experimental conditions (72 h at 32 °C), different results were found in the passerine birds Chinese bulbuls (*Pycnonotus sinensis*) by Wu et al (2014). They found a significant decrease in the body mass after 21 days of heat stress treatment (10 °C vs 30 °C). Groups of quail exposed to 12°C and 33°C showed a percentage body mass loss within 4 days of fasting treatment (Ben-Hamo et al, 2013), which would indicate that a short exposure time to heat stress is not enough to affect the body mass. In chickens, a decrease in weight gain has been seen after 24 hours of exposure to high temperatures (Yahav and Hurwitz, 1996), as in the review carried out by Kumar et al (2021), the weight gain of broilers decreases under high temperatures. But, it can affect some organs, as occurred in our work, where a decrease in intestinal mass was observed in individuals exposed to heat stress. This has also been observed in passerine birds exposed to 28 days of heat stress (Wu et al, 2014). On the other hand, other passerine birds (larks) exposed to temperatures of 35 °C for three weeks decreases their intestinal and stomach mass compared to birds acclimatized to 15 °C (Tieleman et al, 2003). Similarly, chickens under heat stress conditions show decreased jejunal mass, suggesting that high temperatures further reduce intestinal mass and depress enterocyte proliferation and growth (Garriga et al, 2006; Uni et al 2001, Sughiarto et al, 2017). Khaleel et al (2021) observed that broilers exposed to different temperature conditions before hatching, had different values in the length of the villi to depth of the jejunum crypts ratio (VH:CD) in conditions of cold or heat stress for more than 4 days. These data demonstrate that variation in temperature changes intestine morphology. In the case of high temperature, the intestinal mass was lower in the birds, even though the feeding has been *ad libitum* and the few differences may be due to the fact that our studies worked with smaller passerine birds and it is a different species. Despite this, body mass remained unchanged, these observations would indicate that some organs may be more sensitive to heat stress than others.

Regarding hematological parameters, hematocrit was not affected by chronic heat stress. Similar studies in chickens under chronic heat stress confirmed the same pattern (Sughiarto et al, 2017) while acute heat stress exposition (few hours) show a decrease of hematocrit (Yahay & Hurwitz 1996, Altan et al 2003). In our study we found an increase of H/L ratio by heat stress similar to others studies in poultry and wild life (Prieto and Campo 2010; Altan et al, 2003, Altan et al, 2000, Xie et al 2017), in agreement with stress situations and high levels of corticosterone in blood. Several authors have confirmed these data by observing that, when administering corticosterone, the values of the H/L index increased in individuals under stress (Gross et al, 1983; Padrones, 2014, Chediack et al, 2022, Padrones et al, 2023).

It has been observed that some blood parameters such as glucose remain unchanged during heat stress (Sughiarto et al, 2017; Bueno et al, 2017), while others such as triglycerides are modified, in poultry and wild birds (Bogin et al, 1996, Xie et al, 2015; John and George, 1977). In this study, we found an increase in uric acid concentration in plasma, probably by the CORT released during stress. We noticed similar results in house sparrows by injecting CORT (Chediack et al, 2022) and recent evidences show that CORT is probably involved in this process because it induces muscle proteolysis (Furukawa et al, 2021). Moreover, Cohen et al. (2008) proposed that uric acid (UA) may be related to plasma antioxidant capacity rather than protein catabolism. Therefore, they suggest UA as the main circulating molecular antioxidant.

### Capsaicin effect

Industry often use different strategies to lessening thermal stress in poultry, like feed additives. Supplementation of food with feed additives (probiotics, active substances of plants, vitamins, and minerals) increase the heat stress tolerance capacity of chickens (Goel 2021). Capsaicin is a feed additive used often to alleviate stress in poultry (El-Hack et al 2022, Sahin et al 2017, Prieto and Campo, 2010). In alignment with these authors, we notice a decrease in the H/L ratio in house sparrows exposed under heat stress when we orally gavage capsaicin. This give the first evidence of the effects of using this additive in the wild birds. Uric acid shows a decrease after capsaicin administration but not enough to mitigate the thermal stress.

### Digestive physiology: intestinal enzymes

It has been seen in birds that environmental temperature influences both the morphology of the gastrointestinal tract and its digestive functions. Particularly in cold conditions, small birds increase their energy consumption, which in turn compromises their digestive efficiency, associated with an increase in enzyme activity (Karasov et al, 2011). However, to the best of our knowledge a pioneering study of heat stress (acute and chronic) exposition of birds was published by Osman & Tanios (1983) in chickens. In this study, the data shows a biphasic effect of heat stress. Acute heat stress produces an increase in the activity of intestinal maltase and pancreatic amylase, while during chronic stress the activity of digestive enzymes remains stable (Osman & Tanios 1983). Others studies shown the same pattern in digestive and pancreatic enzymes in broilers under acute heat stress (Routman et al, 2003). In our study, we find in house sparrows a similar pattern on the activity of the brush border enzymes (aminopeptidase, maltase and sucrase) and pancreatic enzymes (trypsin and chymotrypsin) under these conditions of chronic heat stress. The activities found in the different portions of the intestine has a classic activity profile (Chediack et al, 2012). The activity of aminopeptidase increasing towards the distal portion, while maltase and sucrose activities has higher values in the proximal and medial portion of the intestine under normal conditions.

These data show that intestinal enzymes do not undergo changes due to heat stress for 3 days (chronic stress), probably because temperature is a regulatory factor for the expression and/or activity of these enzymes when stress is generated in short periods of time.

Regarding the effect of food additives in the diet, it has been observed that in poultry, capsaicin can act as a growth promoter, since a tendency since to increase antibodies has been seen as the capsaicin dose is increased in the diet (Sanabria et al, 2013). Also, as a regulator of normal temperature in chickens with fever induced by LPS (Mahmoud et al, 2007), decreases the heterophil:lymphocyte ratio (Prieto and Campo, 2010), improved jejunal development (width and surface area of the villi) and increased enzymatic activity (lipase and trypsin) (Li et al, 2022). Meanwhile, in rats it has been observed that this compound (the main active compound of the chili pepper seasoning) has different effects “*in vivo*” and “*in vitro*”. “*In vitro*” studies show an increase in the activity of digestive enzymes such as lipase and pancreatic amylase, without affect the activity of chymotrypsin and intestinal disaccharidases (Ramakrishna R. et al, 2003). While, “*in vivo*”, Parkash and Srinivasan (2011) observed an increase in digestive enzymes, pancreatic lipase, amylase, trypsin, and chymotrypsin in rats on a high-fat diet after 8 weeks of capsaicin administration. In our “in vivo” study, we could not find a modulating effect of capsaicin, probably because the time of exposure to capsaicin was not sufficient to generate a significant stimulus to modulate the activity of intestinal enzymes.

In conclusion, a difference of 10 °C in temperature for three consecutive days’ produces heat stress in sparrows, which does not cause significant changes in their body mass, nor in their hematocrit values. However, it modifies some hematological parameters, increases the H/L index and the UA level in plasma, showing that birds are under heat stress. No modification in the activity of digestive enzymes is observed, despite a decrease in the intestine mass. The administration of capsaicin, in sparrows exposed and not exposed to heat stress, does not exert significant effects on body mass or cause changes in hematocrit values. But, capsaicin decreases the H/L ratio in those birds exposed to heat stress. Capsaicin did not produce any modulation of the activity on the digestive enzymes, nor on the intestinal mass.

## Acknowledgments

We appreciate language revision by Dra. Clara Berdasco (University of California, Riverside. USA). This study was supported by UNSL-CyT 02-0820 to FDC-JGC and PICT- 2016-0595 to FDC.

## Notes

**Funding** The research was supported by Universidad Nacional de San Luis N° 02-0820 to F.D.C-J.G.C and Agencia Nacional de Promoción Científica y Tecnológica PICT- 201-0595 to F.D.C.

**Competing interests** The authors declare no competing or financial interests.

### Competing Interest Statement

The authors have declared no competing interest.

